# Attentional failures after sleep deprivation represent moments of cerebrospinal fluid flow

**DOI:** 10.1101/2024.11.15.623271

**Authors:** Zinong Yang, Stephanie D. Williams, Ewa Beldzik, Stephanie Anakwe, Emilia Schimmelpfennig, Laura D. Lewis

## Abstract

Sleep deprivation rapidly disrupts cognitive function, and in the long term contributes to neurological disease. Why sleep deprivation has such profound effects on cognition is not well understood. Here, we use simultaneous fast fMRI-EEG to test how sleep deprivation modulates cognitive, neural, and fluid dynamics in the human brain. We demonstrate that after sleep deprivation, sleep-like pulsatile cerebrospinal fluid (CSF) flow events intrude into the awake state. CSF flow is coupled to attentional function, with high flow during attentional impairment. Furthermore, CSF flow is tightly orchestrated in a series of brain-body changes including broadband neuronal shifts, pupil constriction, and altered systemic physiology, pointing to a coupled system of fluid dynamics and neuromodulatory state. The timing of these dynamics is consistent with a vascular mechanism regulated by neuromodulatory state, in which CSF begins to flow outward when attention fails, and flow reverses when attention recovers. The attentional costs of sleep deprivation may thus reflect an irrepressible need for neuronal rest periods and widespread pulsatile fluid flow.

## Introduction

Sleep plays a fundamental role in maintaining brain health and preserving cognitive performance. Despite the intense biological need for sleep, sleep deprivation is a major facet of modern life. A single night of lost sleep can cause noticeable cognitive impairment, including attentional failures, in which individuals fail to properly react to an easily detected external stimulus (*1–5*). The behavioral deficits caused by sleep deprivation carry a major cost for organismal survival: for example, a momentary lapse in attention while driving a car can have life-threatening outcomes (*6*). Sleep deprivation reliably induces attentional failures despite this cost, suggesting that these deficits reflect an unavoidable need of the brain for sleep. However, the neural basis of sleep-deprivation-induced attentional failures is not yet well understood.

Acute sleep deprivation has widespread effects on both local and global aspects of neurophysiology. Global fluctuations in blood-oxygenation-level-dependent (BOLD) fMRI signals appear across various networks following sleep deprivation (*7*, *8*), and large-scale fMRI signals are linked to altered electrophysiology, eyelid closures (*9*), and arousal state, suggesting global changes in brain activity and vigilance (*10–13*). Attentional lapses after sleep deprivation are linked to reduced activity within thalamus and cognitive control areas, suggesting a failure to engage large-scale network activity (*14–16*). At the scale of local cortical areas, attentional failures are linked to sleep-like low-frequency waves in isolated cortical patches in rats (*17*). Intriguingly, transient increases in low-frequency activity also predict attentional lapses in humans (*18*, *19*). Together, these findings have shown that attentional deficits following sleep deprivation are linked to brainwide hemodynamic changes and low-frequency neural oscillations. A key open question is what drives the brain to generate these spontaneous drops in neural arousal state and behavior after sleep deprivation.

One possibility is that attentional lapses correspond to a brief state in which the brain transiently carries out a sleep-dependent function that is incompatible with waking behavior. Sleep serves many functional purposes, and several studies have shown that one critical role is to clear the neurotoxic waste products that accumulate during wakefulness (*20–24*); although a recent study reported the opposite pattern (*25*). Importantly, the conflicting studies used different sleep protocols for the awake measurements (rested wakefulness vs. sleep-deprived), raising the question of whether sleep deprivation could alter this process. Waste clearance in the brain is mediated by cerebrospinal fluid (CSF), and CSF begins to pulse in large, low-frequency (0.01-0.1 Hz) waves during non-rapid eye movement sleep (NREM) (*26*, *27*). In addition, sleep deprivation modulates pulsatility and solute concentration in the CSF (*28*, *29*), suggesting the effects of sleep deprivation could be accompanied by altered CSF dynamics. However, whether behavioral deficits are linked to CSF flow has not been explored.

We conducted a within-subject total sleep deprivation study in healthy human participants to investigate whether sleep deprivation and its associated attentional deficits are linked to altered brain fluid dynamics. We used multimodal fast fMRI, EEG, behavioral assessments, and pupillometry, to track multiple aspects of neurophysiological dynamics simultaneously. We discovered that CSF flow is coupled to behavioral deficits during wakefulness after sleep deprivation. Notably, we found that attentional failures are marked by a brain and bodywide state change that underlies both behavioral deficits and pulsatile fluid flow, and that the moments where attention fails signal the occurrence of a large-scale pulse of CSF flow in the brain.

## Results

To investigate how sleep deprivation modulates neural activity and CSF dynamics, we performed a simultaneous EEG and fast fMRI experiment in 26 human participants. To enable within-subject comparisons, each participant was scanned twice: once after a night of regular sleep (well-rested), and once after one night of total sleep deprivation, which was continuously supervised in the laboratory (Fig. 1A). We performed simultaneous EEG-fMRI scans with pupillometry in the morning. During the scans, subjects performed up to four runs of a sustained attention task, the psychomotor vigilance test (PVT, Fig. 1B), followed by an eyes-closed resting-state run.

**Fig. 1.**
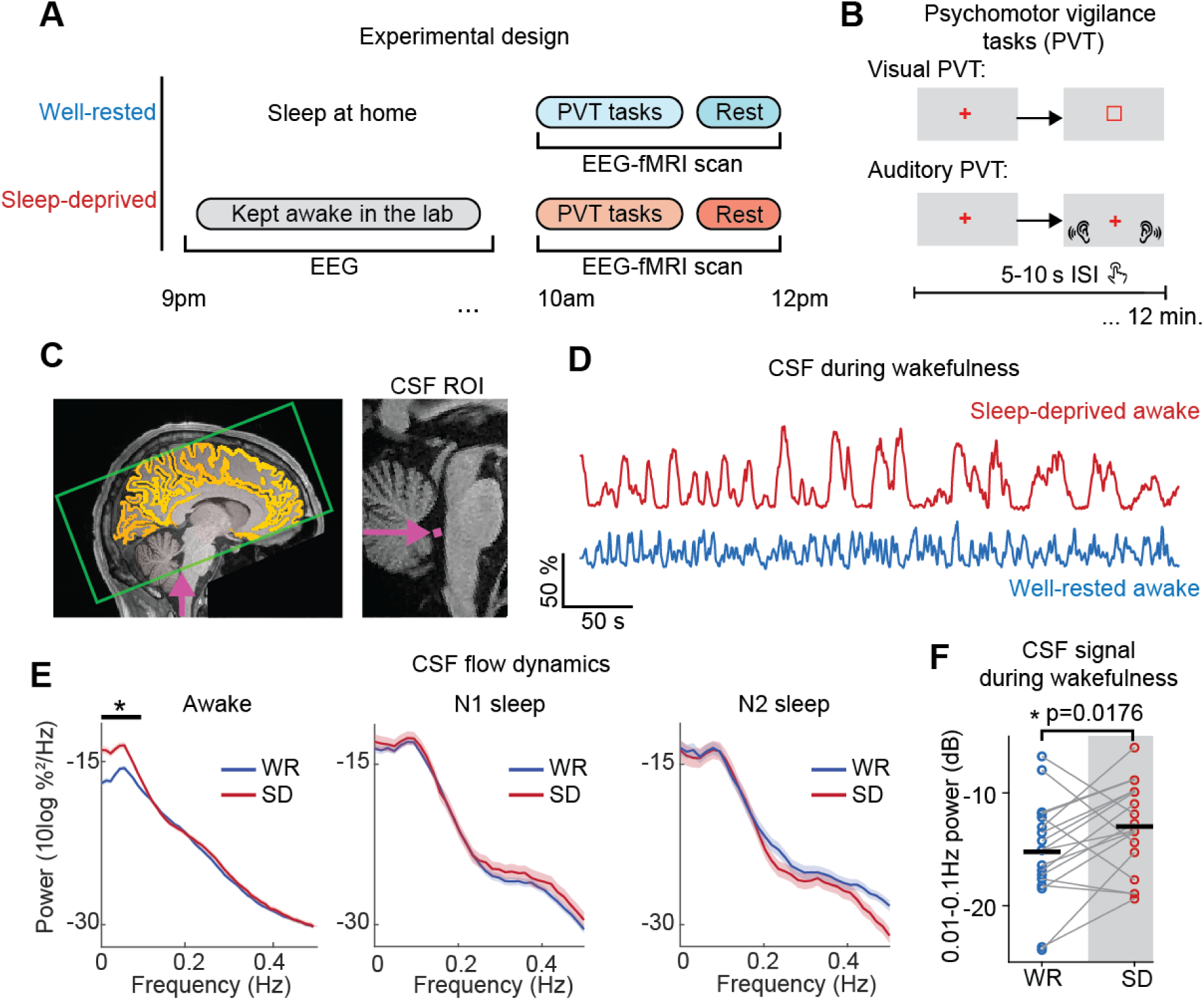
After sleep deprivation, CSF flow exhibits large sleep-like low-frequency waves during wakefulness. (**A**) Sleep-deprived (SD) visit: Subjects arrived at the laboratory at 7PM prior to the sleep-deprived night, and were monitored continuously during the night. Well-rested (WR) visit: Subjects arrived between 8:30-9am the day of the scan. For both visits, scans were performed around 10am in the morning. Scans included up to four PVT runs followed by a 25-minute resting-state run. (**B**) The PVT attention task: Each run used either the auditory PVT (detecting a beep) or the visual PVT (detecting a luminance-matched visual stimulus). **(C**) Left: Example fMRI acquisition volume position (green box) for simultaneous measurement of BOLD and CSF flow. The volume intersects the fourth ventricle, to enable upwards CSF flow detection. Yellow marks the cortical segmentation; purple indicates flow measurement ROI. Image masked to delete identifiers. Right: Example placement of CSF ROI (magenta) in the fourth ventricle in one representative subject. (**D**) CSF timeseries from the same subject during wakefulness shows that sleep deprivation causes large CSF waves during wakefulness, whereas the well-rested (WR) condition shows smaller CSF flow. (**E**) Sleep deprivation increased low frequency (0.01-0.1 Hz) CSF power during wakefulness (Awake n=486 segments in WR; 205 SD; N1 n=179 WR; n=59 SD; N2 n=57 WR; n=40 SD), black bar indicates p<0.05, permutation test with Bonferroni correction), to a magnitude similar to N1 and N2 sleep. Power spectral density (PSD) calculated on CSF signal in resting-state runs. Shading is standard error. (**F**) Paired analysis of CSF low-frequency power in resting-state wakefulness shows increased power after SD (n=18 subjects with artifact-free wakefulness at both sessions, paired t-test, Bonferroni corrected).

### Sleep deprivation causes sleep-like pulsatile CSF flow to intrude into wakefulness

We first investigated whether sleep deprivation altered the dynamics of CSF flow in resting-state scans that included awake and sleeping periods (Fig. S1). Our fast fMRI protocol enabled us to simultaneously measure cortical hemodynamic BOLD signals and CSF inflow in the fourth ventricle (Fig. 1C). Consistent with previous studies, the CSF inflow signal during well-rested wakefulness exhibited a small-amplitude rhythm synchronized to respiration (*30*, *31*), in contrast to NREM sleep where CSF exhibits large ∼0.05 Hz waves (*26*). However, visual inspection clearly showed that the CSF signal during wakefulness after sleep deprivation also exhibited large-amplitude low-frequency waves, resembling NREM sleep (Fig. 1D). We analyzed this CSF signal across sleep stages in participants who had at least 60 seconds of continuous wakefulness during the resting-state run (n=22), excluding segments with high motion (framewise displacement>0.5mm), and found a 4.7 dB increase in the CSF signal peaking at 0.04 Hz in sleep-deprived wakefulness (Fig. 1E). Remarkably, the CSF power in sleep-deprived wakefulness reached levels similar to the CSF power during typical N2 sleep (0.01-0.1 Hz power in SD-wake: −14.96 dB, 95%CI=[−15.88, −14.04]); rested N2: −15.13 dB, 95%CI=[−16.37, −13.88]). Hemodynamics are a key driver of CSF flow, as changes in blood vessels can mechanically drive CSF flow (*26*, *32–34*), and we found that sleep deprivation also induced a significant increase in low-frequency (0.01-0.1 Hz) cortical gray matter BOLD power (p=0.0048, paired t-test). These session-dependent changes in CSF and BOLD were not driven by motion artifacts (Fig. S2). These results demonstrated that sleep deprivation caused low-frequency CSF flow pulsations and hemodynamic waves to appear during wakefulness, resembling the dynamics typically seen during N1 or N2 sleep.

### Pulsatile CSF flow occurs during epochs with worse attentional task performance

Since large-scale CSF waves typically occur during NREM sleep, a state in which attention is suppressed, we investigated whether the CSF waves during sleep-deprived wakefulness were associated with any attentional cost. As expected (*2*, *14*, *35–41*), sleep deprivation caused an increase in the mean reaction time (Fig. 2A), and the omission (missed response) rate (Fig. 2B) in the PVT task. Surprisingly, we observed that attentional lapses (RT>500ms) and omissions tended to coincide with higher-amplitude CSF flow (Fig. 2C). To quantify this behavior-CSF relationship, we tested whether lapses (RT>500ms) and omissions were linked to CSF power, analyzing data in 60-s segments during confirmed wakefulness (EEG and no eyelid closures>1s). We found that worse attentional performance was linked to increased low-frequency (0.01-0.1 Hz) CSF power: segments with attentional lapses and segments with omissions exhibited significantly higher CSF flow power than segments without lapses (Fig. 2D). This effect remained significant when controlling for motion (two-way ANOVA; p>0.05 for motion). These results demonstrated that larger low-frequency CSF flow is associated with attentional failures during wakefulness.

**Fig. 2.**
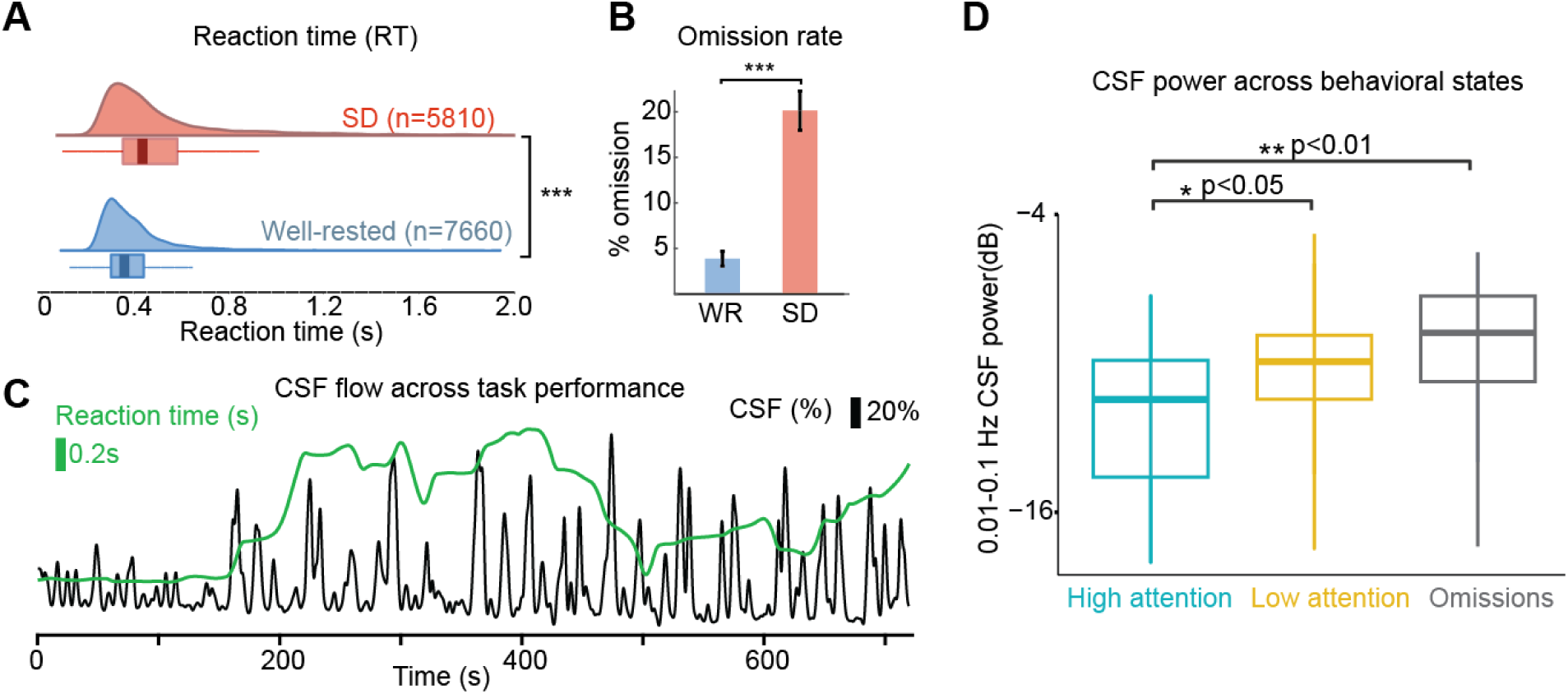
Pulsatile CSF flow dynamics increase during epochs with slower reaction times and attentional failures. (**A**) Reaction times (RTs) after sleep deprivation showed higher mean and a longer tail, indicating more behavioral lapses (Mann-Whitney U test, p<0.001). (**B**) Omission rate increased after sleep deprivation (n=26 subjects, p<0.001, paired t-test). (**C**) CSF flow and reaction time fluctuations during one example run, showing higher flow when reaction time slows down. (**D**) Low-frequency (0.01-0.1 Hz) CSF power within non-overlapping 60s segments, categorized into three different states: high attention (all RTs below 500 ms); low attention (at least one RT>500 ms), and omissions (at least one omission). Higher CSF power appears in lower attentional states (p<0.001 for main effect; one-way repeated measures ANOVA and Tukey post hoc test, n=26 subjects).

What process could link CSF flow to attentional function? Ascending neuromodulation, such as noradrenergic input from the locus coeruleus, modulates attentional function (*42*, *43*), and also act directly on the vasculature (*44*), making this a candidate mechanism for modulating both attention and CSF flow. Neuromodulatory tone and locus coeruleus firing is correlated with pupil diameter (*45*), so we investigated whether pupil diameter was linked to CSF flow on faster timescales. We found that pupil diameter was significantly correlated with CSF flow: constricted pupil was linked to downwards CSF signals, and dilated pupil was linked to subsequent upwards CSF signals (Fig. 3 A&B). This correlation remained significant when we tested segments from rested wakefulness and after sleep deprivation (Fig. S3). Consistent with this, we also observed that CSF flow peaks were locked to drops in attentional performance: both reaction time and the rate of omissions increased before CSF peaks (Fig. 3 C&D). In each case, while CSF flow was locked to behavior and pupil, it lagged these measures in time, which could reflect a vascular mechanism which would introduce a time delay. Furthermore, while pupil diameter is consistently correlated with noradrenergic tone, it is also correlated with several other modulators (*46*, *47*). To test whether pupil-CSF coupling reflected a pupil-locked modulation of cerebrovascular fluctuations, which could be expected from noradrenergic-driven vasoconstriction and would drive a delayed CSF response (*48*), we also examined the global cortical BOLD signal, and found an anticorrelation with the pupil (Fig. 3E). We calculated the best-fit impulse response linking pupil to the global BOLD signal, and found a negative impulse response with a peak delay of 6.75s, consistent with pupil-linked vasoconstriction. To test whether this mechanism in turn predicted CSF flow, we convolved the pupil signal with its impulse response and calculated the expected CSF flow driven by this vascular effect, with no additional parameter fitting (Fultz et al., 2019). We found that the convolved pupil signal yielded a significant prediction of CSF activity (Fig. 3H, zero-lag R=0.26, maximal R=0.3 at lag −1.75s), indicating that the timing of the pupil-CSF coupling could be explained by a vascular intermediary.

**Fig. 3.**
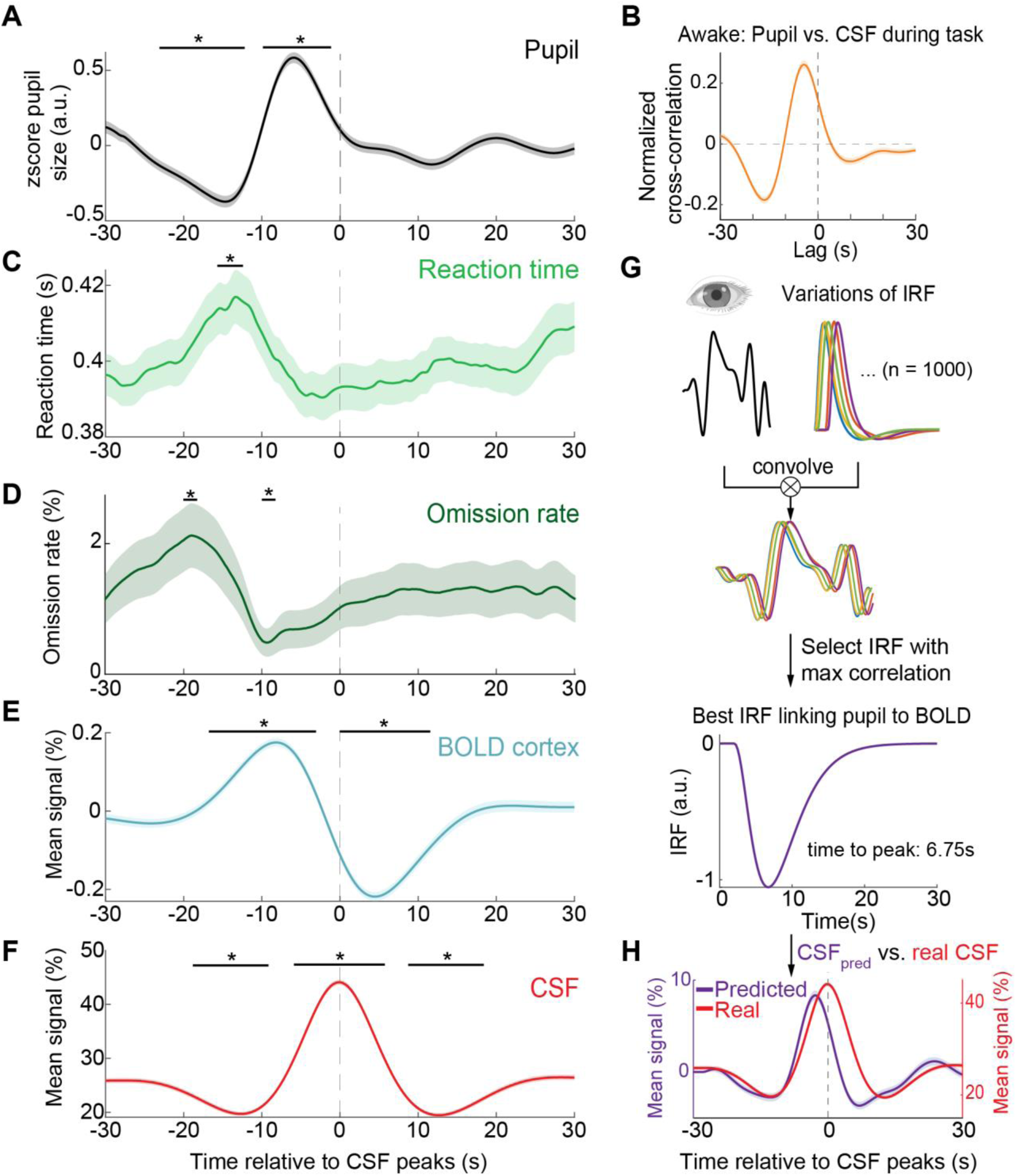
Pulsatile CSF flow is temporally coupled to pupil diameter changes and behavioral performance during wakefulness. (**A**) Spontaneous pupil constriction and dilation is time locked to peaks of CSF flow. Black bars indicate significant changes in zscored pupil diameter compared to baseline (p<0.05, t-test, baseline = [−30 −28] s, Bonferroni corrected). (**B**) Cross correlation between pupil diameter and CSF showed strong correlation (maximal r = 0.26 at lag −4.25s; n = 709 segments, 26 participants) (**C**) During period with pupil constriction, reaction times also showed significant increase compared to same baseline (p<0.05, t-test, Bonferroni corrected). (**D**) Omission rate during task showed significant increase during pupil constriction, and significant decrease during pupil dilation (p<0.05, t-test, Bonferroni corrected). (**E**) Significant biphasic changes in cortical BOLD activity are locked to CSF peaks (p<0.05, t-test, Bonferroni corrected). (**F**) Mean CSF signal. (**G**) To estimate the impulse response function linking pupil size changes to BOLD and CSF activity, we convolved pupil diameter traces from each segments with a series of impulse response function (IRF). Estimated impulse response of the cortical BOLD signal to the pupil diameter shows a time-to-peak at 6.75s. Cross-correlation between pupil diameter and BOLD showed strong correlation (maximal r = 0.38 at lag 0s; n = 709 segments, 26 participants). (**H**) Predicting CSF flow with no additional parameter fitting, assuming that the derivative of the pupil-locked BOLD fluctuations drives CSF flow, shows significant prediction of the true CSF signals (zero lag r = 0.26; n=709 segments).

### CSF pulsatile flow is temporally coupled to attentional failures and altered brain state

Having found that CSF flow was highest in epochs with omissions, we next examined the precise dynamics occurring at the moment of omissions. Since CSF flow increases during NREM sleep, we carefully excluded omissions due to NREM sleep, to confirm whether these flow changes appeared during wakefulness (EEG-verified, and excluding segments with eye closures>1s). We found a striking coupling between behavior and CSF flow during sleep-deprived wakefulness: omissions were locked to a downward wave and then an upward wave of CSF flow in the fourth ventricle (Fig. 4A). While few omissions occurred in the well-rested state, when they did appear they showed similar CSF flow coupling, demonstrating that a complete attentional failure even when rested can engage a similar mechanism (Fig. S4A). The cortical gray matter BOLD signal exhibited a matched biphasic pattern, consistent with vascular drive of downwards and then upwards CSF flow (Fig. S4B). However, our imaging protocol was only able to directly measure CSF inflow (upwards to the brain) due to the placement of the acquisition volume (Fig. 1C), whereas drops in CSF signal could in theory represent either outflow or simply no flow. We therefore conducted a second study in an additional 10 sleep-restricted subjects with a new imaging protocol that measured bidirectional flow (Fig. S6A) to test whether this pattern specifically signaled CSF outflow before inflow. We confirmed that omissions were locked to a pulse of CSF flowing downward out of the brain, beginning at the time of the missed stimulus, followed by a pulse flowing upward into the brain (Fig. S6B). These results replicated the finding of coupled attentional function and CSF flow in an independent cohort, with a biphasic profile of downwards flow at omissions, followed by upwards flow.

**Fig. 4.**
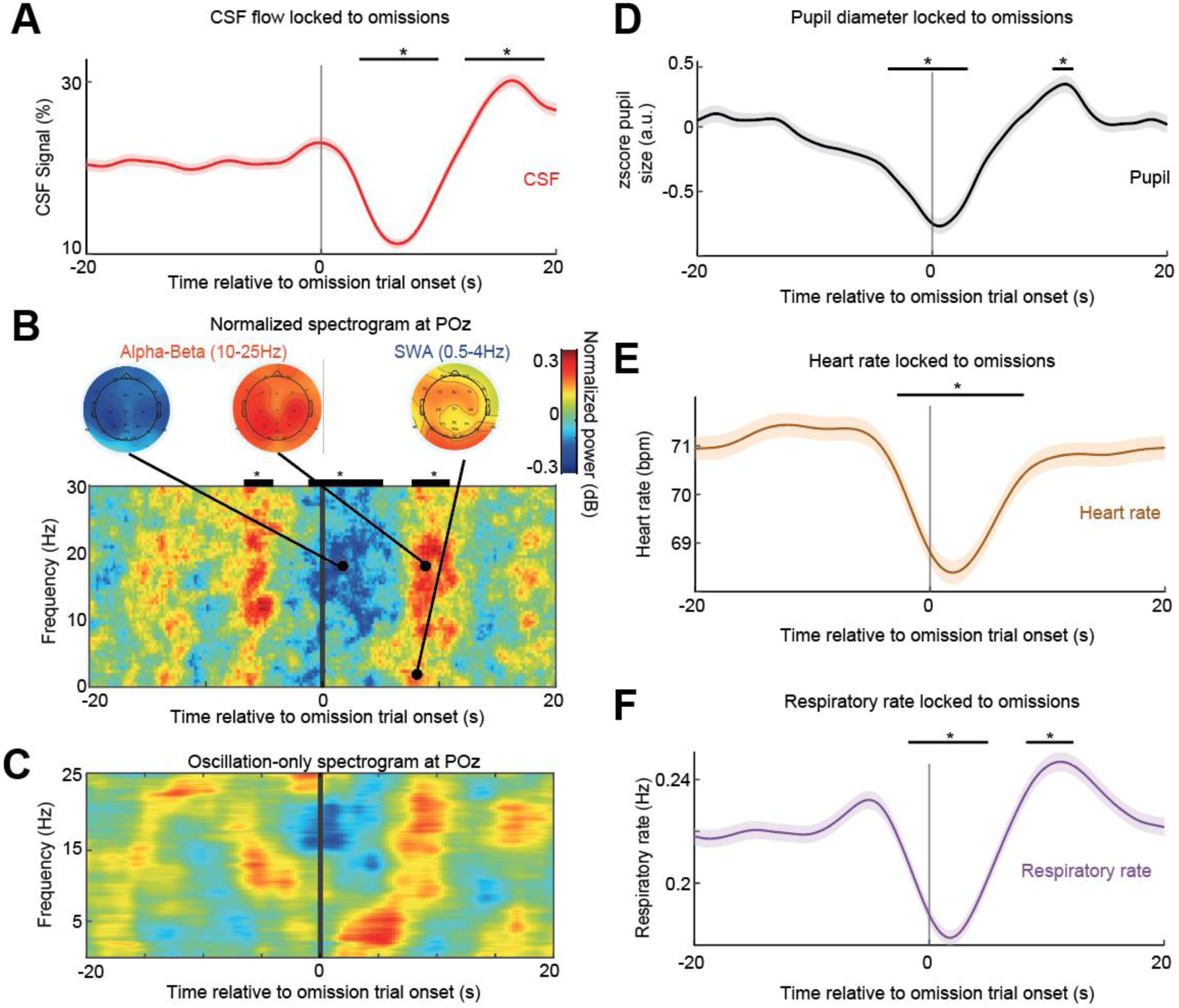
Attentional failures are coupled to pulsatile CSF flow and a series of neural and systemic physiological changes. (**A**) Awake omission trials are locked to a biphasic change in CSF flow, with a downwards trough followed by an upwards peak. Time zero marks stimulus onset; omission occurs within the following seconds. (**B**) Omission trials are locked to a broadband drop in EEG power, including decreased alpha-beta (10-25Hz) EEG power then an increase in slow wave activity (SWA; 0.5-4 Hz) and alpha-beta power. The biphasic alpha-beta power change is widespread with centro-occipital predominance, whereas the SWA increase has occipital and frontal predominance. Spectrogram is normalized within-frequency; black bars indicate significant a change in broadband power (0.5-30Hz) compared to baseline (p<0.05, t-test, baseline=[−20 −10] s, Bonferroni corrected). (**C**) The aperiodic component of the EEG was subtracted to display oscillation-specific changes. The alpha-beta and SWA effects were still present (statistics in Fig. S5). (**D**) Pupil diameter showed a biphasic change at omissions, with a significant constriction during the omission trial followed by dilation after trial onset (9.00-12.00s). (**E**) Heart rate dropped during omissions and subsequently increased. (**F**) Respiratory rate dropped at omissions and subsequently increased. Black bars with stars indicate significant changes from baseline (n=364 trials, 26 subjects, p<0.05, paired t-test, Bonferroni corrected). Shading is standard error.

During NREM sleep, CSF flow is coupled to neural slow wave activity (*21*, *25*), and local slow waves can also increase after sleep deprivation (*17*). We therefore analyzed the EEG spectrogram to investigate whether neural dynamics might be linked to these omission-locked CSF waves. We found that omissions (n=364 trials, 26 subjects) were accompanied by EEG alterations across multiple frequency bands, with a drop in broadband EEG power, particularly pronounced in the alpha-beta (10-25Hz) range, followed shortly by subsequent increase in power (Fig. 4B). In addition to this broadband change, the oscillation-only component of the spectrogram (calculated by subtracting the aperiodic component (*49–52*) also demonstrated significant clusters at slow and alpha-beta frequencies, with a broad spatial distribution (Fig. 4C, Fig S5). Overall, this broadband power reduction and steepening spectral slope pointed to a spatially distributed change in electrophysiological dynamics.

What type of neuronal event might these EEG spectral shifts reflect? The relatively broadband nature of these EEG changes suggested that they might reflect a transient change in cortical excitability (*53*), perhaps mediated by central neuromodulatory state, as suggested by the pupil coupling (Fig. 3). In addition, autonomic state changes and systemic oscillations can drive CSF flow (*54–56*), consistent with the idea that large-scale neuromodulatory changes could contribute to both the attentional failures and CSF dynamics. We therefore tested whether a sequence of events could underlie the CSF flow: a drop in attentional state followed by recover. Consistent with this, pupil dilation was biphasically-coupled to CSF flow: the pupil was constricted during the omission, corresponding to a low arousal state and downward CSF flow, and then subsequently dilated (Fig. 4D), with CSF then flowing upward (Fig. 4A). Since neuromodulatory state also has systemic effects (*57*), we further examined systemic physiological recordings, including changes in respiratory rate and volume, heart rate, and peripheral vascular volume – reflecting activity of the autonomic nervous system. We found that all systemic measures were significantly locked to the omission (Fig. 4D-F, Fig. S7). These results demonstrate that CSF inflow events during wakefulness were coupled to a brain-and-bodywide integrated state shift, manifesting as a transient behavioral deficit, electrophysiological markers of distinct neuronal state, and pupil constriction.

### CSF pulsatile flow is differentially modulated by loss and recovery of attention

These results clearly identified widespread behavioral, neuronal, and arousal dynamics that were locked to CSF pulsatile flow during sleep-deprived wakefulness. However, attentional failures are typically brief during wakefulness, with a spike in arousal rapidly following an omission. Due to this biphasic pattern, this CSF-locked result could be explained by two different possibilities: either that CSF flow was specifically linked to drops in arousal (at the onset of attentional failures) or that it was locked to increases in arousal (at recovery of behavior). We therefore designed an analysis of behavioral responses to separate the neurophysiological dynamics linked to the drop, vs. recovery, of attention. We categorized behavioral omissions into three types: “Type A”, where the omission trial was preceded and followed by a valid response; “Type B”, the first of at least three consecutive omissions; and “Type C”, the last omission of a consecutive series (Fig. 5A). As above, wakefulness during all omissions was verified both through EEG scoring and eye-tracking.

**Fig. 5.**
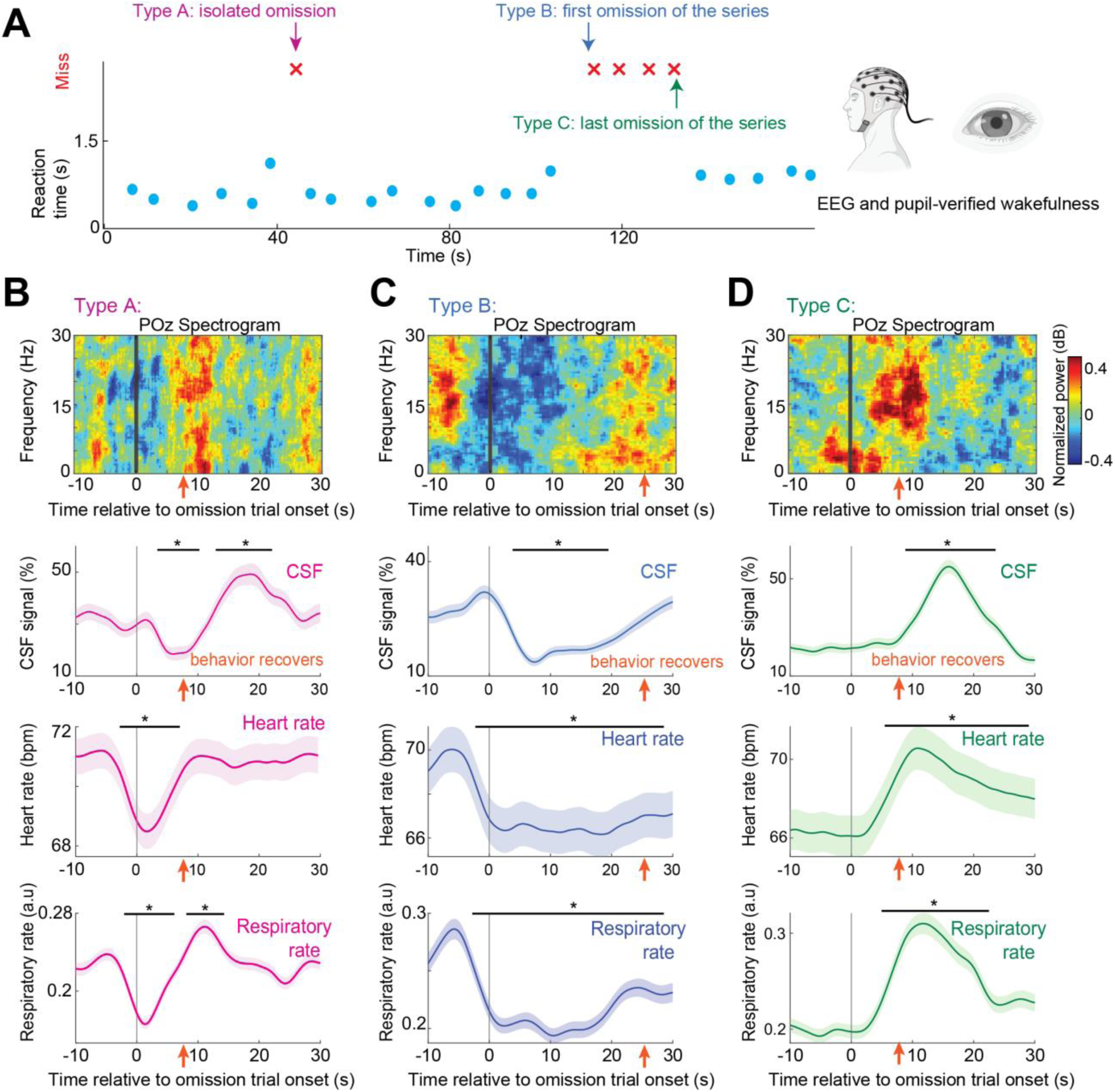
CSF flows outwards when attention drops, and back in when attention recovers. (**A**) Omissions during wakefulness were categorized as isolated (Type A); onset of sustained low attention (Type B); and end of sustained low attention (Type C). Schematic made with BioRender. (**B**) Type A, isolated omission trials (n=127 trials, 26 subjects) were locked to a decrease in broadband EEG power, then increased power at attention recovery in the next trial. CSF showed a significant decrease followed by a significant increase. Respiratory rate and heart rate both showed significant changes from baseline after trial onset. Orange arrow points to the average timing (8.1s) of the first valid response after the isolated omission. (**C**) Type B: first omission of the series signifies the onset of a sustained low attentional state (n=61 trials, 20 subjects). CSF signal, respiratory rate and heart rate all decrease significantly (p<0.05, paired t-test, Bonferroni corrected). Orange arrow points to the average timing (25.1s) of the first valid response after the series of omissions. (**D**) Type C: last omission of the series, followed by increased attentional state (n=57 trials, 20 subjects). EEG shows heightened SWA during the omission and subsequent increase in alpha-beta power at recovery of attention. CSF signal, respiratory rate and heart rate all increase significantly (p<0.05, paired t-test, Bonferroni corrected). Orange arrow points to the average timing (9.0s after time 0) of the first valid response after the series of omission. For all panels in B-D: Black bars indicate significant (p<0.05, paired t-test, Bonferroni corrected) changes from baseline ([−10 −5] s). Shading is standard error.

Remarkably, this analysis identified two separable patterns: neural, CSF and systemic signals showed opposing changes locked to Type B (arousal drops) and C (arousal increases) omissions (Fig. 5B, C, D). The loss of attentional focus in Type B omissions was linked to a decrease in EEG alpha-beta power and expulsion of CSF, whereas regaining of attentional focus in Type C omissions co-occurred with an increase in EEG broadband power and upwards flow of CSF, and the systemic arousal indicators followed the same pattern (Fig 5C, D). These results demonstrated that CSF flow occurs in a specific sequence of events: during an attentional failure, EEG power dropped in concert with bodywide signatures of low arousal, and a pulse of CSF flows downwards out of the brain. This pattern inverted at recovery of arousal, with CSF returning upwards as attention improved, with a time delay consistent with vascular-mediated CSF flow. These results identified widespread behavioral, neuronal, and arousal dynamics that were locked to CSF pulsatile flow during sleep-deprived wakefulness, consistent with an integrated neuromodulatory system governing neural arousal state and CSF flow.

## Discussion

Here, we found that attentional failures after sleep deprivation are locked to a brain- and body-wide state change and pulsatile waves of CSF flow. During sleep-deprived wakefulness, CSF flow pulsations intruded into the awake state, causing a flow pattern that resembled N2 NREM sleep. Strikingly, CSF waves were strongly coupled to behavioral failures, a shift in neuronal spectra, and pupil constriction. Specifically, attention dropped when CSF flowed outward, and attention recovered as CSF was drawn back upwards into the brain. The attentional failures that occur after sleep deprivation thus reflect the initiation of a process in which CSF is transiently flushed out of the brain.

Our data showed a consistent pattern of broadband EEG shifts linked to attentional failures. What type of neural event might these represent? The process of falling asleep is linked to suppression of EEG alpha power and drops in respiratory rate (*58*, *59*), which could suggest that the events we observe here represent an accelerated transition toward sleep onset. In addition, autonomic arousal fluctuations are characterized by a sequential modulation of beta power and slow wave activity (*54*, *60*, *61*). The EEG pattern we observe here shares similar broadband features, and we find that it represents a behaviorally relevant modulation of neural vigilance state, with attentional consequences. This pattern may thus reflect that sleep deprivation causes sleep-initiating cortical events to occur during wakefulness, but with an interruption of the initiation process before sleep is attained. Given that we found a shift in spectral slope, this pattern could also be consistent with increasing cortical inhibition (*62*). While we could not directly identify local sleep slow waves here due to the macroscopic nature of these measurements, this effect appeared to be relatively spatially widespread, as it was detected across many EEG electrodes.

The neural and attentional drops were also linked to pupil constriction, a classic signature of arousal state that is actively controlled by noradrenergic neuromodulatory projections (*63*, *64*). Pupil diameter is correlated with attention, heart rate, and galvanic skin reflex, reflecting a tight coupling between the state of the central and peripheral arousal systems (*12*, *47*, *65*). Furthermore, the autonomic system can strongly modulate CSF flow, with strong effects during light sleep (*55*). The high correlation between pupil diameter and CSF pulsatile flow thus suggests the involvement of the ascending neuromodulatory system in regulating CSF flow during wakefulness (*66–68*). A possible mechanistic explanation for the joint changes in EEG, CSF, and autonomic indicators is the well-established parallel projections from key neuromodulatory nuclei such as the locus coeruleus, which projects throughout cortex and thalamus to modulate neural state, while also directly acting on the sympathetic nervous system (*69–72*). Intriguingly, locus coeruleus activity oscillates during NREM sleep and drives infraslow neuronal spectral changes (*73*, *74*), and pupil diameter also covaries with these neural signatures (*75*, *76*); our results suggest a similar dynamic of temporally structured noradrenergic fluctuations could appear during sleep-deprived wakefulness. Other neuromodulators are also correlated with pupil diameter and could contribute (*46*, *77*), although our hemodynamic data suggest a substantial role for the vasoconstrictive effects of the noradrenergic system (*78*). All of the dynamics we report were observed during eyes-open wakefulness, demonstrating that brief shifts in attentional state are sufficient to elicit these coordinated changes. Our results could thus be consistent with sleep deprivation causing a destabilization of the ascending arousal system, with brief failures of its wakefulness-promoting actions leading to attentional dysfunction, neuronal arousal suppression, and CSF flow.

A key question is why these attentional shifts produce such large-scale CSF flow, detectable even in the ventricle. The neuromodulatory mechanism could potentially explain these dual effects. A primary driver of CSF flow is changes in the vasculature, as dilation and constriction of blood vessels in turn propels CSF flow (*26*, *32*, *33*, *79–83*). Since noradrenaline is a vasoconstrictor (*44*), drops in locus coeruleus activity could decrease neuronal arousal state while dilating cortical blood vessels and constricting the pupil. This would result in a downwards flow in the fourth ventricle detected several seconds later, which then would invert as attention recovers and noradrenergic activity increases, reflected in the subsequent pupil dilation. An exciting implication of these dynamics is that pupil diameter could potentially provide a noninvasive, accessible readout of signals related to CSF flow.

These findings may impact the interpretation of previous fMRI experiments. While common practice is often to regress out CSF signals to attempt to remove noise, our data show that this regression-based approach will also remove or distort attention-related signals of neural origin that are collinear with the CSF signals – which are often signals of interest when studying brain dynamics. Moreover, CSF flow signals reported here are within eyes-open wakefulness, suggesting that even during the awake state, fluctuations in attention could have strong effects on CSF signals. Notably, CSF flow signals linked to eye closure have been detected in the widely used Human Connectome Project dataset (*84*), suggesting that regressing out CSF may influence a large set of fMRI studies.

A surprising finding of this study is that attentional function is coupled to CSF flow, with slower reaction times and attentional failures corresponding to the initiation of CSF flow out of the brain. Prolonged wakefulness leads to buildup of toxic metabolic waste products (*22*, *23*, *29*, *85*), suggesting that more fluid flow would be needed after sleep deprivation. Why would behavioral performance and fluid dynamics then be tightly coupled at rapid timescales within the sleep-deprived awake state? One possibility is that they are not directly functionally related, but rather that the same circuit controls both and thus they covary. In this case, drops in ascending arousal could drive both behavioral changes (to decrease waste production) and fluid changes (to increase waste transport). The timing of events we observed could align with this hypothesis, since the downward CSF flow in the fourth ventricle began at the onset of the missed stimulus, and could reflect a change in the cortex and subarachnoid space seconds earlier. An alternate possibility is that prolonged CSF flow is incompatible with stable attentional states, for example because high flow rates alter the concentration of signaling molecules that are required for typical waking function. We found that while CSF flow was tightly coupled to behavior, it did not exclusively occur during omissions − CSF flow pulses continuously with lower amplitude during wakefulness (*30*, *56*, *86*), and flow increased to a lesser degree during successful performance with slow reaction times (Fig. 2, 3) − meaning that a single large event of CSF flow does not prevent behavior. However, it is possible that sustained high-amplitude flow over longer timescales interferes with behavioral performance, leading to a structured alternation between high-attention and high-flow states.

Overall, our results demonstrate that the attentional failures caused by sleep deprivation are coupled to large-scale brain fluid transport. When the brain lacks the opportunity to sleep, it enters a sub-optimal attentional state, which may provide partial benefits of sleep at the cost of behavioral errors. This coupling points to a central circuit that controls both attentional state and CSF flow, and shows that the moments of attentional dysfunction we experience after sleep loss correspond to the emergence of widespread fluid flow in the brain.

## Acknowledgments

We thank B. Tan, B. Dormes, J. Licata, M. Bosli, M. Aon, Z. Valdiviezo, I. Vinal, N. Tacugue, N. Leonard, T. Ly, Z. Diamandis, D. Zimmerman, J. Yee, M. Ruiz, J. Hua, R. Huang for assisting with data collection, S. Chakrapani and S. McMains for MRI support. This research was funded by National Institutes of Health grants U19NS128613, R01AT011429, R00MH111748, and R01AG070135, an NDSEG Graduate Research Fellowship to S.D.W., and a NAWA Fellowship to E.B., and the McKnight Scholar Award, Sloan Fellowship, Pew Biomedical Scholar Award, One Mind Rising Star Award, and the Simons Collaboration on Plasticity in the Aging Brain (#811231). This work used resources provided by NSF instrumentation grant 1625552.

## Competing interests

LDL is an inventor on a pending patent application for an MRI method for measuring CSF flow.

## Supplementary Materials

### Materials and Methods

#### Participants

The main experiment enrolled 32 healthy subjects, of whom 26 subjects successfully completed both study visits (7 males; 19 females; mean age=25.6 years±4.41 years, range=19 to 40 years). All 26 subjects completed two experimental sessions spaced 8 to 10 days apart: a well-rested session, and a sleep-deprived session. For the well-rested (WR) condition, subjects completed a full night of sleep at home, and they arrived at the lab no later than 9am for a 10am scan start time. In the sleep deprivation condition (SD), subjects arrived at the lab at approximately 7pm and stayed overnight until completing the scan the following day. Subjects were screened to ensure they had no history of sleep, medical or psychiatric disorders, and good typical sleep quality (sleep schedule of 6.5-9h/night), as assessed by self-rating questionnaires. Exclusion criteria included shift workers, subjects that had traveled across time zones in the 3 months before the study, and any MR contraindication. All subjects were required to maintain a regular sleep schedule for one week before the visit, with sleep prior to scans monitored by wrist actigraphy, to consume less than 100 mg caffeine daily prior to the study, and to refrain from alcohol, caffeine, and sleep-aid medications for 24 hours prior to the day of the scan. All subjects provided written informed consent and study procedures were completed as approved by the Boston University Charles River Campus Institutional Review Board (IRB #5059E).

#### Overnight experimental design

During the SD visit, subjects were sleep deprived overnight under constant monitoring by two staff members. At least two researchers remained in the adjacent control room to verify continuous wakefulness during SD. Upon arrival, subjects were fitted with an EEG cap for overnight monitoring. Subjects wore eye-tracking glasses (Tobii Pro 3 Glasses) throughout the night except when performing testing sessions, for continuous eye monitoring while the staff members logged any spontaneous long eyelid closures throughout the night. Research staff continuously monitored the live video of participant’s eye from the control room during the sleep deprived night. If the participant closed their eyes for more than two seconds, a staff member requested that they open their eyes and then engaged them in conversation to ensure wakefulness. If the eye closure repeated frequently, staff members took subjects on a gentle walk. Between 6AM and the morning scan, subjects took a shower, during which verbal confirmation of wakefulness was obtained, and then they were provided with breakfast.

EEG was recorded during four task testing blocks (TB) during the night of sleep deprivation (TB1: around 21:00, TB2: around 23:00, TB3: around 2:00, TB4: around 5:00). Each block consisted of three eyes-open resting state runs (2 minutes) and two vigilance task runs (12 minutes each). The vigilance task consisted of 12-minute runs of a computerized auditory psychomotor vigilance task (aPVT) and visual PVT (vPVT, details in next section). Auditory and visual runs were counterbalanced in order. When not involved in testing sessions, subjects were allowed to carry out their own preferred activities, such as reading, writing, listening to music, watching TV or playing games. Lying down, sleeping and vigorous physical activity were not permitted. Non-scheduled light snacks were permitted, while caffeinated beverages, alcohol and medications that can influence sleepiness were not allowed during the deprivation protocol.

#### Visual and Auditory PVT

Analyses of PVT included the data from both the aPVT and vPVT tasks. During the aPVT (adapted from (*1*, *2*)), subjects were instructed to respond as fast as possible (by pressing a button) to a 0.5s 375 Hz tone. Tones were presented binaurally at a comfortable volume through MR-compatible earbuds (Etymotic Research, Inc., Elk Grove Village, Illinois, ER3 transducers). Cushions were placed beside ears to restrict movement and to further reduce scanner noise. During the vPVT, subjects were instructed to respond as fast as possible (by pressing a button) when the fixation crosshair changed into a square. The two visual stimuli were designed to have identical luminance to avoid light-induced pupillary responses. For both task modalities, each stimulus lasted 0.5s and inter-trial intervals ranged from 5-10 s. A fixation crosshair was presented throughout. The total duration of each task run was 12 min. Visual stimuli inside the scanner were presented on a VPixx Technologies PROPixx Lite Projector (VPixx Technologies, Quebec, Canada) with a refresh rate of 120Hz.

#### MRI experimental design

Subjects underwent five fMRI runs: two runs of aPVT (12 minutes each), two runs of vPVT (12 minutes each) and one run of eyes-closed resting state (25 minutes). Auditory and visual run order were counterbalanced. 16 subjects successfully completed all four task runs for both visits; 10 subjects completed a subset of these runs (e.g. due to inability to stay awake in the later runs). If subjects did not successfully complete all task runs, the successfully completed runs were still included in the analysis. Runs were excluded if subjects failed to respond to more than 20 trials, indicating they could not stay awake. This resulted in 198 runs from 26 subjects (6 runs were excluded). During the eyes-closed resting state session, subjects were instructed to press a button on every breath in and breath out as long as they were awake and were told that it was permitted to fall asleep during the scan.

#### Sleep behavior analysis

Sleep-wake behavior in the week before the SD visit were collected by actigraphy and stored as the sum (activity) of 30 s intervals. These data were analyzed with Actilife (v.6.7, Actigraph, Pensacola, FL), using a sleep/wake detection validated algorithm. Subjects were also instructed to keep notes of bed and rise times to help frame the time in bed during which actigraphy data were analyzed.

### EEG and fMRI data acquisition

Subjects were scanned on a 3 Tesla Siemens Prisma MRI scanner with a 64-channel head/neck coil. MRI sessions began with a 1mm isotropic multi-echo MPRAGE anatomical scan. Functional scans consisted of single-shot gradient echo multi-band EPI sequences with 40 interleaved slices (2.5mm isotropic resolution, TR=378ms, MultiBand factor=8, TE=31 ms, flip angle=37, FOV=230x230, blipped CAIPI shift=FOV/4, and no in-plane acceleration).

Additional sensors were used to record systemic physiology during the MRI scan: respiration was measured simultaneously using an MRI-safe pneumatic respiration transducer belt around the abdomen and pulse was measured with a photoplethysmogram (PPG) transducer (BIOPAC Systems, Goleta, California, USA). Physiological signals were acquired at 2,000 Hz using Acqknowledge software and were aligned with MRI data using triggers sent by the MRI scanner.

EEG was collected using a 64-channel MR-compatible EEG cap and BrainAmp MR amplifiers (Brain Products GmbH, Gilching, Germany). Two re-usable reference layers were custom-made to record ballistocardiogram artifacts (BCG), as in (*3*). As described in (*3*), with the use of a reference layer, a subset of EEG electrodes can directly record the BCG noise, enabling effective modeling and removal of this noise. This artifact signal can then be used to regress out the BCG artifact from the EEG signals. The first reference layer, used for subjects 1-6, was made from an ethylene vinyl acetate shower cap (the inner, insulating layer) and a layer of modal-lycra fabric (the outer layer), which were fastened together by round, plastic grommets to create openings large enough to accommodate the EEG electrodes, as in (*4*). These grommets were placed over electrodes Fpz, Fz, AFz, FCz, Cz, Pz, POz, Oz, F3, F4, F7, F8, T7, T8, C3, C4, P3, P4, PO3 and PO4, and held in place with cyanoacrylate glue. Holes were cut in the reference layer to site each grommet, and larger holes were also made as to allow the subjects’ ears to pass through. The outer layer of the reference layer was moistened with a solution of baby shampoo and water and potassium chloride to make it conductive prior to each recording. Finally, the inner face of the EEG cap was lined with Nexcare surgical tape (3M, St. Paul, Minnesota), to avoid wetting the EEG cap. The second reference layer, used for subjects 7-26, was designed as reported in (*3*). It was made from a layer of stretch vinyl (Spandex World Inc., Long Island City, USA) coated in PEDOT ESD500 and P1900 ink (PEDOTinks.com, Austin, USA), with grommets placed over Fp1, Fp2, F3, F4, C3, C4, P3, P4, O1, O2, F7, F8, T7, T8, P7, P8, Fpz, AFz, Fz, FCz, Cz, Pz, POz, and Oz, and large holes for the subjects’ ears. For this reference layer, no conductive solution or tape was needed. Prior to each recording, electrode impedances were measured. To ensure safety, subjects were not allowed to proceed to EEG-fMRI recordings unless the impedance of all channels was below 100 KΩ; however, a lower impedance of less than 50 KΩ was targeted for signal quality. EEG was collected at a sampling rate of 5000Hz, synchronized to the scanner clock, referenced to channel FCz.

### EEG preprocessing

Gradient artifacts were removed through average artifact subtraction (*5*), using a moving average of the previous 30 TRs. All electrodes were then re-referenced to the common average, separately for electrodes contacting the head and the electrodes placed on the reference layer. EMG and ECG channels placed on the chin and at the back were excluded from the common average. Ballistocardiogram artifacts were removed using regression of reference electrode signals (*4*). Because the position of electrodes and physiological noise dynamics can vary over the recording sessions, we implemented a dynamic time-varying regression of the reference signals in sliding windows (*6*, *7*).

### EEG spectral analysis

We used EEGLAB (*8*) and MATLAB (R2022a, MathWorks Inc., Natick, MA) to analyze EEG data. Continuous EEG data were downsampled to 500 Hz and then bandpass filtered with EEGLAB anti-aliasing filter to 0.2–40 Hz. Independent component analysis (ICA) was used to remove components reflecting eye movements and muscle activity. EEG power spectra density (PSD) was calculated using multitaper spectral estimation (Chronux (*9*)), using 5s windows and 5 tapers. Further analysis was performed on EEG channel POz since this channel consistently had good quality data across subjects. EEG POz channel was used to calculate PSD with 5s windows and 5 tapers. For normalized EEG power spectrogram analysis, power values for each trial and each frequency were z-scored.

Analyses of the spectral slope of the EEG signal collected over the night of sleep deprivation (outside of fMRI scanner) used channel POz. The power spectral density (PSD) from 0.5-35 Hz was estimated using the multitaper method in 10s epochs with 0.1s overlap which were subsequently used to extract the slope of a fitted model using the FOOOF algorithm in the 1 to 35Hz range (*35*). To separate the aperiodic and oscillatory component of the PSD, we used FOOOF to calculate an initial linear fit of the spectrum in log-log space, and subtracted the results from the spectrum. Then a Gaussian function was fitted to the largest peak of the flattened PSD exceeding two times the SD. This procedure was iterated for the next largest peak after subtracting the previous peak until no peaks were exceeding the minimum peak threshold. FOOOF settings were kept default with the exception of minimum peak height limits set to be over 0.3 dB. The oscillatory components were finally obtained by fitting a multivariate Gaussian to all extracted peaks simultaneously. After the iterations, the initial fit was added back to the flattened peak-free PSD, resulting in the aperiodic component of the PSD. Afterward, this aperiodic component was fitted again, leading to the final fit with aperiodic intercept and exponent as parameters. The fitted Gaussian functions were referred to as oscillation-only spectrogram. The aperiodic features were estimated using a sliding window approach with 5s window length and stepping every 0.3s.

### Sleep scoring

One subject in the well-rested session and one subject in the sleep deprived session did not have valid EEG recordings during the resting-state scan and were thus excluded from sleep analysis. The remaining 24 subjects’ sleep EEG data were used in sleep scoring and segmenting the BOLD fMRI data into sleep stages (Fig. 1E,F; Fig. S1). Sleep scoring was performed on preprocessed EEG data using the Visbrain toolbox by two scorers following the standard AASM scoring guidelines (*10*, *11*). Disagreement in sleep scores were further discussed and resolved with a third experienced scorer.

### Eye tracking & pupillometry

Eye tracking over the sleep deprived night (8PM-6AM next day) was done with eyeglasses (Tobii Pro Glasses 3) with rear-facing infrared cameras. Pupil size was recorded in the morning during EEG-fMRI scans using an MR-compatible eye-tracker (EyeLink 1000; SR Research, Osgoode, ON, Canada) at a sampling rate of 1000Hz. The eye-tracker was placed at the end of the scanner bore, such that the participant’s left eye could be tracked via the head coil mirror. Before each task run, we began with a calibration of the eye-tracker using the standard five-point EyeLink calibration procedure. Moments when the eye-tracker received no pupil signal (i.e., during eye blinks) were marked automatically during acquisition by the manufacturer’s blink detection algorithm. Pupil data was preprocessed offline. Missing and invalid data due to blinks were replaced using linear interpolation for the period from 100 ms before blink onset to 400 ms after blink offset. Missing data due to long eyelid closures (>1s) were marked but not interpolated, since these segments were later used to detect microsleeps. Following the automated interpolation procedure, the data were manually checked and corrected if any artifacts had not been successfully removed. 21 runs were excluded from pupil diameter analysis due to technical issues with the eye-tracker, and 6 were excluded due to failed video recording, yielding 171 valid runs from 26 participants. Fig. 3 analyses used data from these 171 runs. We normalized the pupil diameter timeseries to z-scores. For analyses of pupil responses aligned to either CSF peaks or omission trial onsets, we epoched the pupil time series around each event of interest relative to event onset. We rejected pupil epochs if they contained >1s consecutive long eyelid closures within the raw data.

### MRI preprocessing

The cortical surface was reconstructed from the MPRAGE volume using recon-all from Freesurfer version 6 (*12*, *13*). All functional runs were slice-time corrected using FSL version 6 (slicetimer; https://fsl.fmrib.ox.ac.uk/fsl/fslwiki) and motion corrected to the middle frame using AFNI (3dvolreg; https://afni.nimh.nih.gov/). Each motion-corrected run was then registered to the anatomical data using boundary-based registration (bbregister).

We defined the left and right hemisphere cortical gray matter regions of interest (ROIs) using the automatically generated Freesurfer-derived segmentations of cortical gray matter. Cortical time series were converted to percent signal change by dividing by the mean value of the time series after discarding the first 20 volumes to allow the signal to reach steady state.

### CSF flow analysis

CSF inflow analysis used the raw acquired fMRI data with slice-timing correction but without motion correction, as motion correction corrupts the voxel slice position information needed for inflow analysis, and motion correction cannot be accurately performed on edge slices where tissue moves in and out of the imaging volume. All analysis therefore was performed on segments with framewise displacement<0.5, and segments with high motion were marked and rejected from any further analysis. To identify the CSF ROI, we manually traced the intersection of the fourth ventricle with the bottom 3 slices of the imaging frame for each run for each subject. We extracted the mean signal in this region as the CSF inflow signal. All CSF inflow time series were highpass filtered above 0.01Hz and converted to percent signal change by dividing by the value of the time series the 15th percentile value (since the CSF signal has a floor due to only measuring upwards flow with the acquisition scheme used in Experiment 1). CSF time series were upsampled to a new sampling frequency of 4 Hz using spline interpolation before stimulus-locked averages were calculated.

### CSF peak-locked analysis

Peaks in the band-pass filtered CSF signal (0.01-0.1 Hz) during tasks were identified by identifying local maxima that surpassed an amplitude threshold of 30%. These peaks (n=709) were then used to extract a peak-locked signal (−30s to 30s relative to the CSF peaks) in pupil diameter (band-pass filtered at 0.01-0.1Hz). To test for statistical significance of the peak in the pupil signal, we used a time-binned, two-sided t-test, averaging the signal in 2-s bins, testing whether the signal was significantly different from baseline ([−30 −28]s in Fig. 3). The p-value was Bonferroni corrected for the number of time bins, and time bins with corrected p-values less than alpha=0.05 were considered significant.

### IRF model estimation

We generated 1000 candidate impulse response functions using the gamma distribution with varying shape and rate parameter. We selected the impulse response function that yielded the highest correlation between the IRF-convolved pupil and the cortical gray matter BOLD signal.

### Spectral power analysis

Spectral power of BOLD and CSF signals was calculated using Chronux multitaper spectral estimation (5 tapers). Power was estimated in the 60 s segments classified as awake, N1, N2, and N3, and the mean power in each participant was calculated from all low-motion (framewise displacement<0.5) segments. N3 was not analyzed because only 5 participants entered N3. Statistical testing across spectra was performed with a permutation test that shuffled segments, and then compared in frequency bins corresponding to the bandwidth (0.05 Hz), with Bonferroni correction for the number of frequency bins. Pairwise comparisons of mean low-frequency power (0.01–0.1Hz) in Fig. 2D,E were computed within the participants who had awake data in both well-rested and sleep deprived visits (n=18) using a paired t-test.

### Behavioral analysis

Behavioral analyses (Fig. 3) included data from both the aPVT and vPVT tasks. Because mean performance varies across visual vs. auditory tasks, the omission rate comparison was performed separately for visual and auditory PVT (Fig. 3B; p<0.001 for each task).

### Omission-locked analysis

For Figure 4, awake omissions (n=364) were identified in 26 subjects, after excluding any omissions with simultaneous eyelid closures longer than 1s, excluding any segments scored as sleep based on the EEG, and excluding high-motion segments. All omissions during confirmed-wakefulness periods were marked and their stimulus onset time were used to extract omission-locked signals in the EEG, pupil diameter, heart rate, respiratory flow rate, respiratory volume per time, peripheral vascular tone and the CSF signal. Analyses of the spectrogram of the EEG signal collected inside the fMRI scanner used channel POz, calculated in the 0.5–30 Hz range using the multitaper method in 10s epochs with 0.1s overlap. They were subsequently used to extract the oscillation-only spectrogram using the FOOOF algorithm in the 1 to 25Hz range. To test for statistical significance of clusters in EEG spectrogram and EEG oscillation-only spectrogram, the spectrograms were averaged and compared to a baseline window spectral power (−20s to −10s relative to omission onset) using a paired two-tailed t-test. The p-values corresponding to each pixel of the spectrogram were corrected with a false discovery rate (FDR) less than 0.05. To test for statistical significance of the peak in the CSF, EEG broadband power, pupil, and systemic physiology signals, we considered the window [−20 −10]s prior to the omission trial onset as baseline. Statistical significance was then calculated for the remaining window in 1-s bins: the signal in each bin was averaged and then compared to baseline using a paired (within-trial) two-sided t-test (p<0.05). The p-value was Bonferroni corrected for the number of time bins tested.

Additionally, three types of omissions (A, B and C) were separately analyzed by finding omissions that were preceded and followed by valid responses (Type A), the first of three or more consecutive omissions (Type B), and the last of three or more consecutive omissions (Type C). Eye-tracking videos were used to identify and exclude trials with long eyelid closures (LEC>1s). In total, 62 series of continuous omissions without LEC were identified. One Type B event and five Type C events were rejected due to high motion. The remaining omissions (Type A n=127, Type B n=61, Type C n=57) were marked and their stimulus onset times were used to extract omission-locked signals. Because this analysis focused on post-trial effects at recovery vs. drops of attention, the analysis window was set to [−10 30]s and baseline set to [−10 −5] s.

### Systemic physiology

The transducer recording the PPG signal was placed on the left index finger, making sure heartbeats were visible in its signal. MRI scanner triggers were co-recorded with the physiological signals to allow data synchronization. We first removed artifactual spikes from the data by replacing time points exceeding 4 standard deviations above the mean with the mean signal level. Then, low-pass filtering was performed on the PPG and respiration belt signals at 30 and 10 Hz cut-off respectively. From the PPG signal, an indicator for peripheral vascular volume (PVV) was derived by calculating the standard deviation of the PPG signal in 3s segments, which is a measure of the excursion amplitude caused by the cardiac beat and is proportional to the volume of blood detected by the PPG sensor (*14*). The variation in HR was computed by averaging the differences between pairs of adjacent heartbeats contained in the 6 s window around each trigger and dividing the result by 60 (beats per minute). The filtered chest belt signal was used to derive a measure of respiratory flow rate, which previously was found strongly to affect the fMRI signal throughout the brain (*15*). This was done by taking the derivative of the low-pass filtered belt signal, rectifying it, and applying a secondary low-pass filter at 0.13 Hz cut-off. The respiratory volume per time was calculated using the standard deviation of the respiratory waveform on a sliding window of 6s centered at each trigger.

### Bidirectional flow experiment

The bidirectional flow measurement in Fig. S3 used an independent cohort and new acquisition protocol designed to simultaneously measure upward and downward CSF flow. 10 healthy adults (4 female, 6 male; age range 22 to 29 years, mean 24.1) were included. The same health and MRI exclusion criteria as for Experiment 1 were used. Subjects participated in a single scan session scheduled for 10AM. To induce attentional failures, participants underwent self-monitored sleep restriction in their homes, sleeping only 4 hours the night before the scan. Scans were performed on a 3T Siemens Prisma scanner with a 64-channel head and neck coil. Anatomical scans used a 1 mm isotropic multi-echo MPRAGE. Functional runs were acquired using TR=593ms, 2.5mm isotropic resolution, 8 slices, FOV=230X230, TE=31ms, no multiband, flip angle=51°, and no in-plane acceleration. The acquisition volume was positioned with the top slice intersecting the narrow opening of the fourth ventricle, and the bottom slice at the base of the ventricle (as in the main experiment), allowing us to image both upwards and downwards CSF simultaneously. Analyses were performed locked to all isolated omissions, with an omission immediately preceded and followed by a hit. For analysis of outflow dynamics, the CSF ROI was manually traced with the top 3 slices of the imaging volume; for the inflow dynamics, the CSF ROI was traced with the bottom 2 slices of the imaging volume. All CSF outflow and inflow time series were converted to percent signal change by dividing by the value of the time series by the 15th percentile value. The percent signal change CSF outflow signal was inverted for display (Fig. S3) so that down represented downward flow in both timeseries.

---

**Fig. S1.**
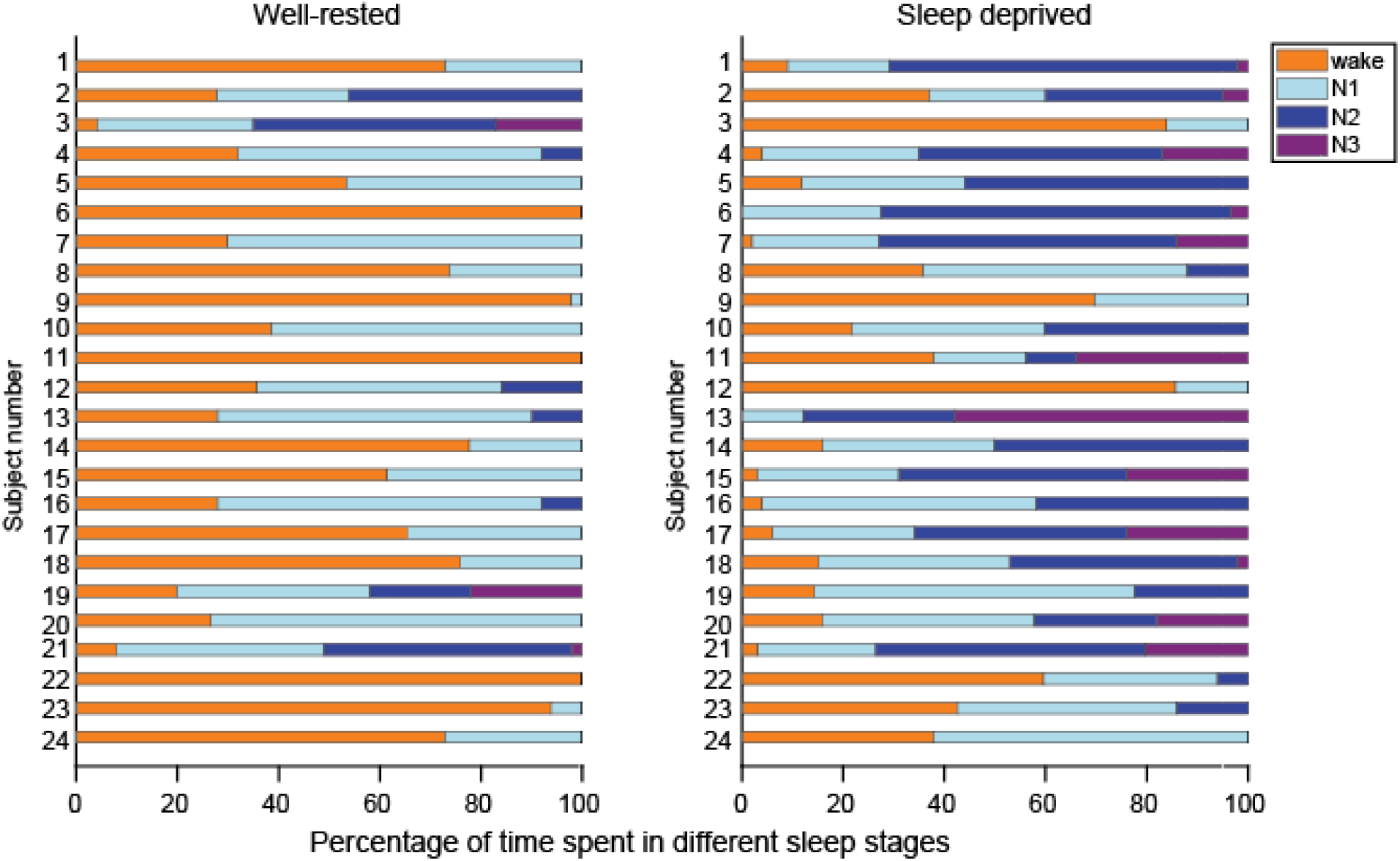
Summary of sleep data. 24 subjects have valid EEG recordings during the rest scan at both well-rested and sleep deprived visits. Their sleep stages are scored according to the current AASM standard.

**Fig. S2.**
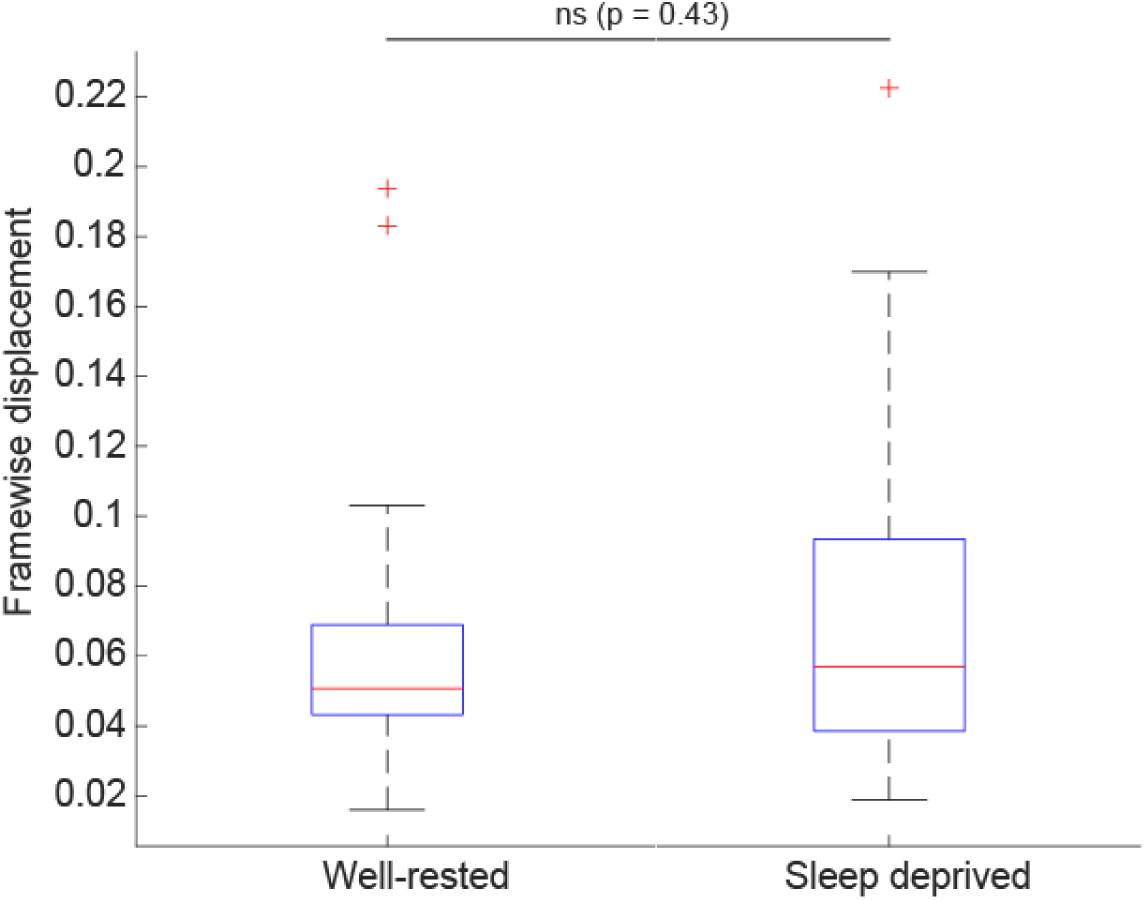
Motion across sessions. Mean framewise displacement (FD) values from the analyzed 60s segments (after excluding frames with >0.5mm FD) used in pairwise comparison in Figure 2 D&E (N=18 subjects). With our motion exclusion criteria there was no significant difference in motion in the analyzed segments across conditions (p=0.43).

**Fig. S3.**
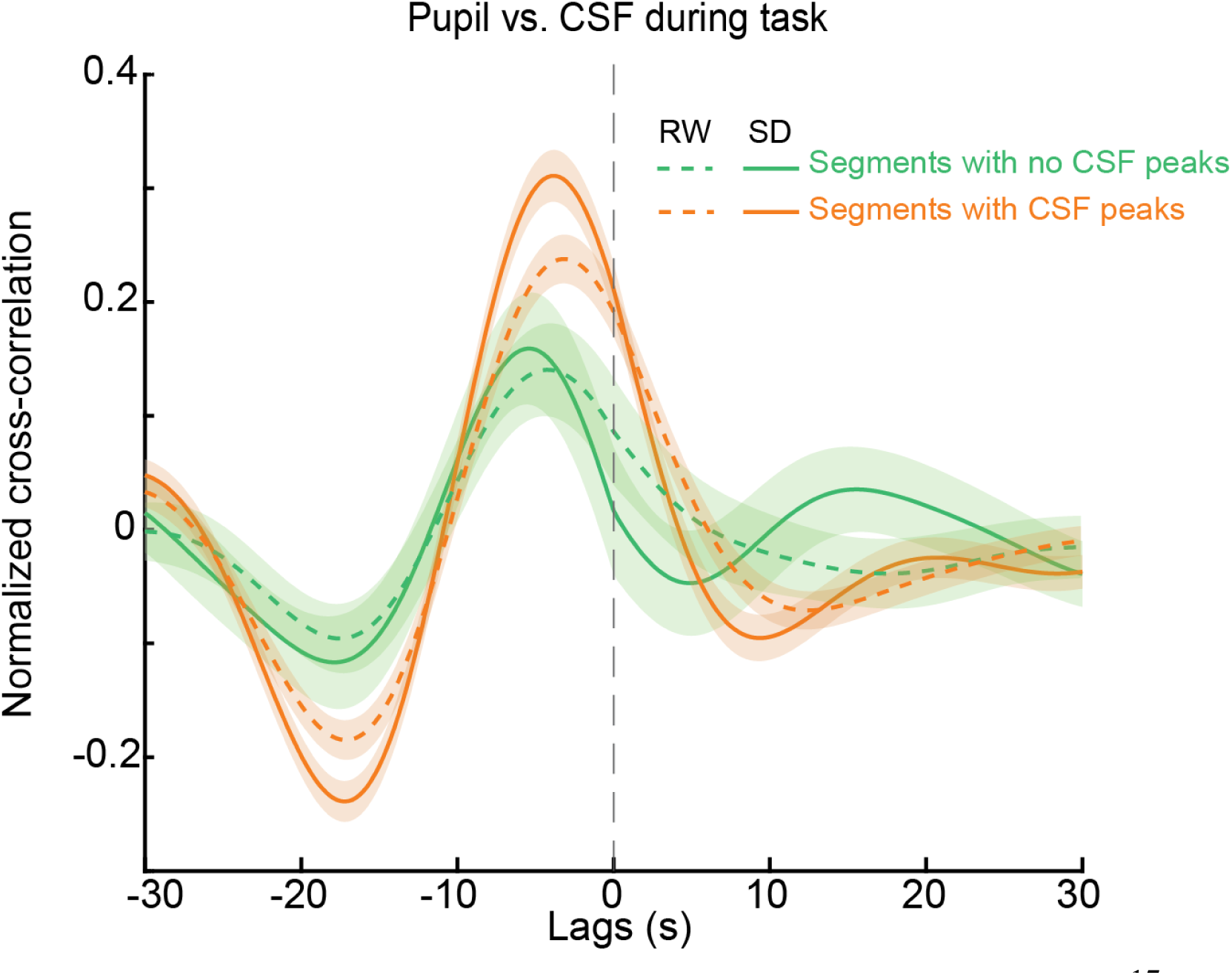
Pupil cross-correlation with the CSF signals during wakefulness. Cross-correlation was separately computed between pupil and CSF during four different conditions: 1) Segments in rested wakefulness with no CSF peaks, 2) segments after sleep derivation with no CSF peaks, 3) segments in rested wakefulness with at least one CSF peak, 4) 60s segments in sleep deprivation with at least one CSF peak. Maximal cross-correlation’s r was observed during SD condition with at least one CSF peak. Segment length was 60 s.

**Fig. S4.**
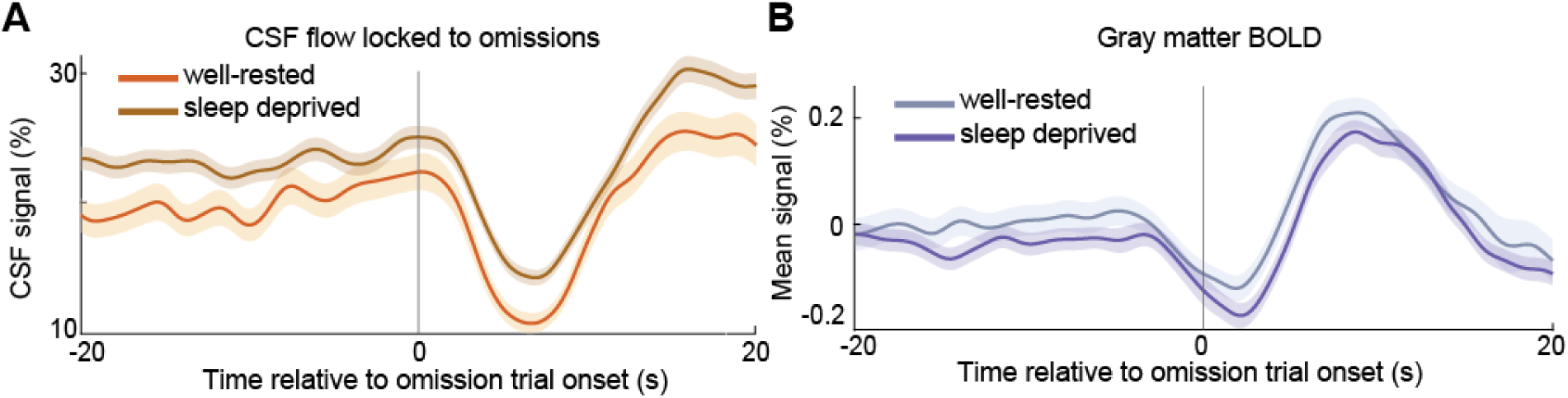
The cortical gray matter BOLD and CSF signal is locked to omissions both during well-rested and sleep-deprived sessions. (**A**) CSF flow at both well-rested and sleep deprived session showed biphasic change that’s locked to the omission trial onset. Mean CSF signal at sleep deprived session is 3.3% higher than the mean CSF signal at well-rested session. (**B**) Cortical BOLD signals in both well-rested (n = 87) and sleep-deprived (n=277) sessions showed a similar biphasic pattern as the ventricular CSF, although with a dip in signal amplitude seconds before the omission trial onset.

**Fig. S5.**
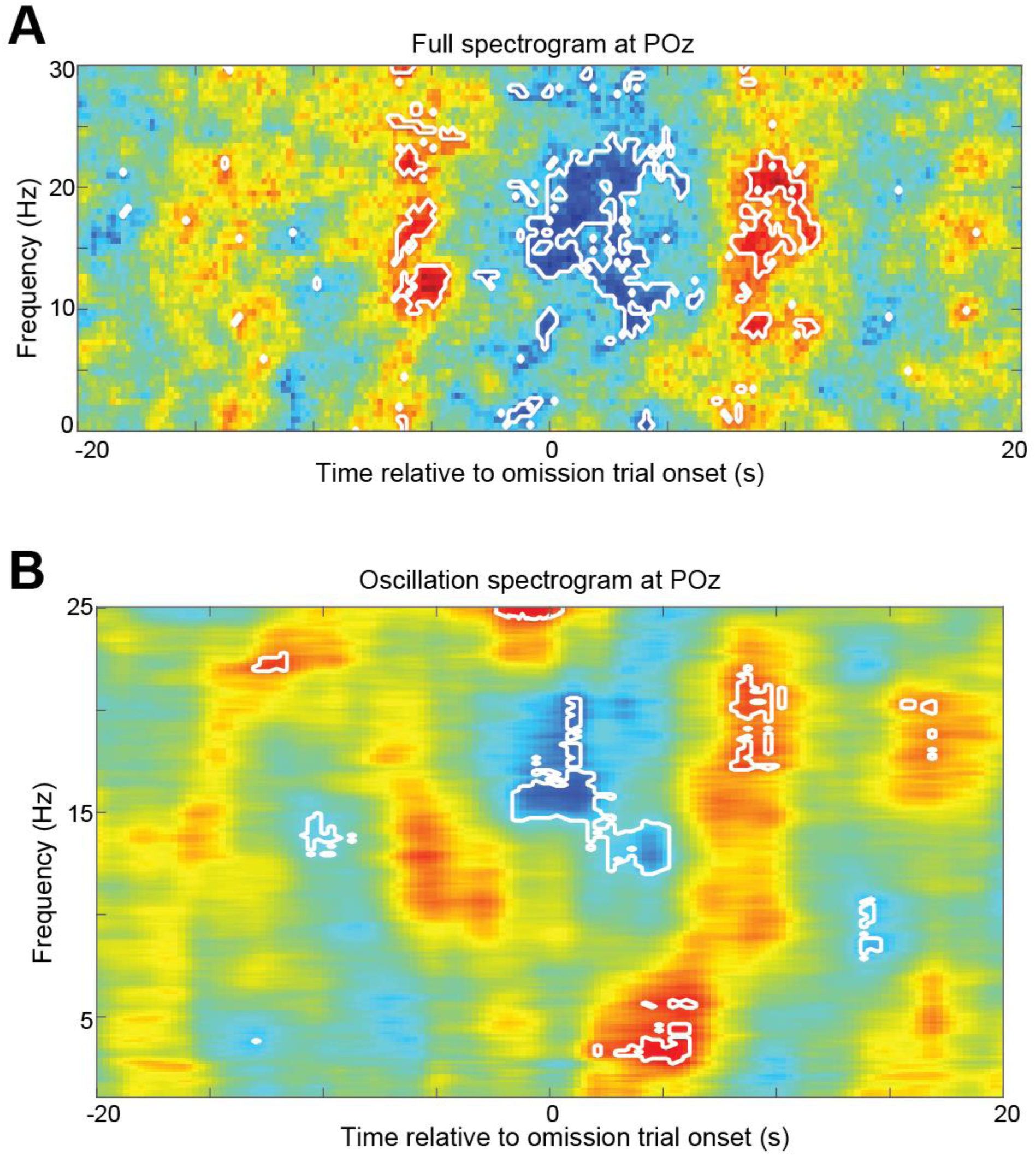
Full and oscillation-only EEG spectrogram at omission trial onset. (**A**) Contour indicates the period of significant power increase with FDR correction p<0.05. (**B**) Contour indicates the period of significant oscillatory activity in SWA and alpha-beta (10-25Hz) range with FDR correction p<0.05.

**Fig. S6.**
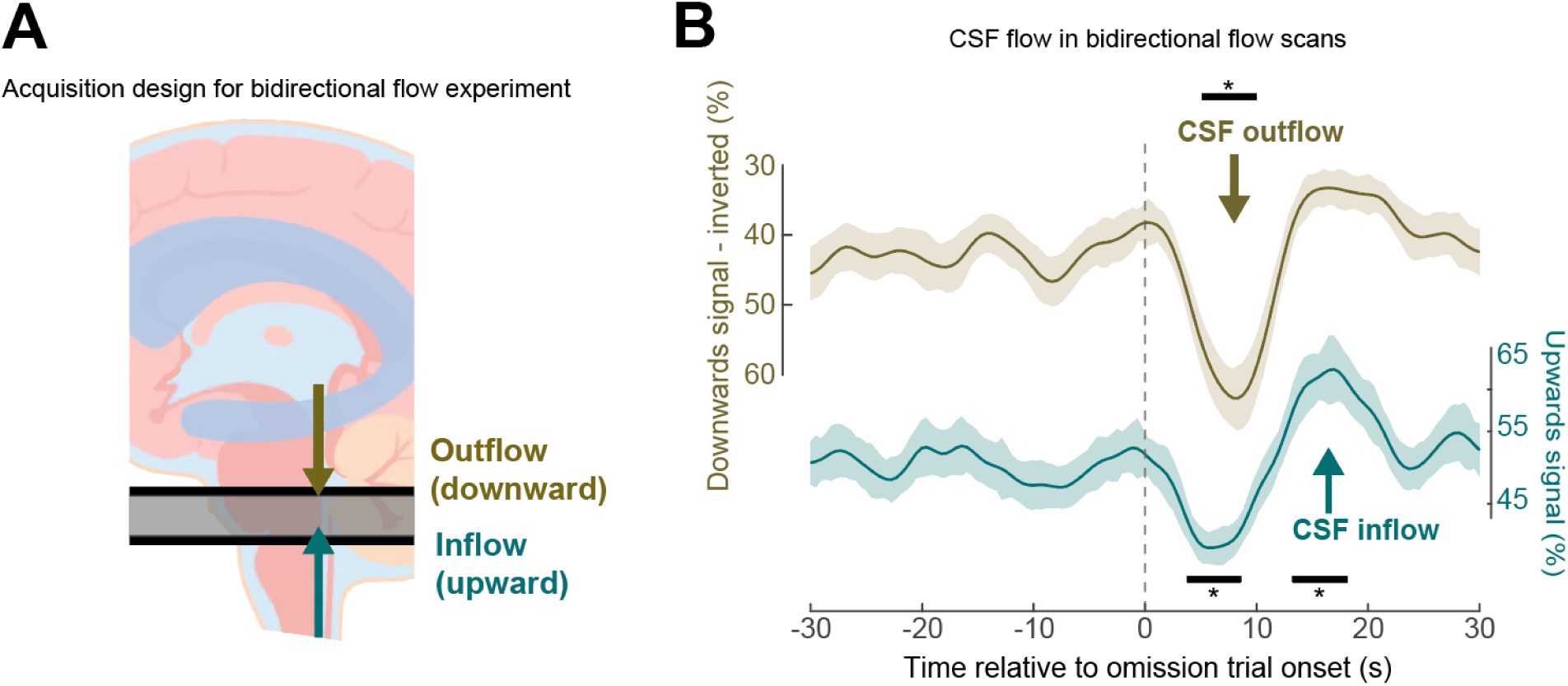
Bidirectional flow imaging experiment demonstrates that omissions are locked to downward then upward CSF flow. **(A)** Positioning of acquisition protocol for the second experiment, designed to measure bidirectional flow. Acquisition volume enabled simultaneous measures of downward flow (measured in the top slice) and upward flow (measured in the bottom slice). (**B)** This bidirectional imaging experiment replicated the coupling between omissions and CSF flow. CSF flow moves downwards at the omission, and moves upward at the recovery of attention. (n=114 trials, 10 subjects). CSF outflow signal is inverted to match directionality (down is downwards flow). CSF flow is averaged relative to isolated omissions during wakefulness. Black bars indicate significant (p<0.05, paired t-test, Bonferroni corrected) changes from baseline ([−10 −5] s). Shading is standard error.

**Figs. S7.**
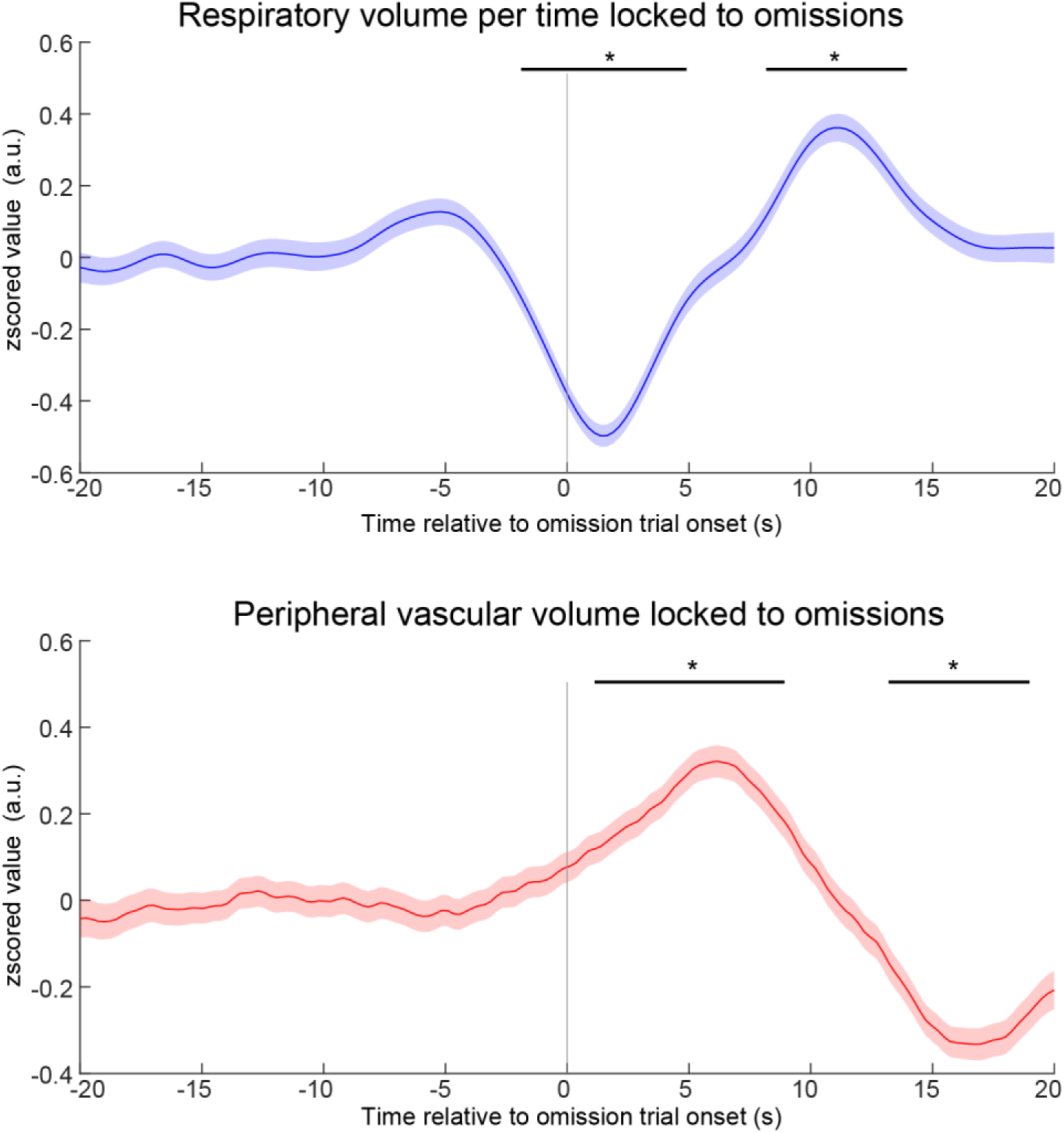
Additional systemic physiology measures are coupled to omission trials. Top: Respiratory volume per time showed biphasic changes locked to omission trials onset. Bottom: Peripheral vascular volume also showed biphasic changes locked to omission trials onset. Black bars indicate significant difference from baseline (paired t-test, Bonferroni corrected, p<0.05); same segments as in Fig. 3.

